# Enrichment and physiological characterization of a novel comammox *Nitrospira* indicates ammonium inhibition of complete nitrification

**DOI:** 10.1101/2020.06.08.136465

**Authors:** Dimitra Sakoula, Hanna Koch, Jeroen Frank, Mike SM Jetten, Maartje AHJ van Kessel, Sebastian Lücker

## Abstract

The recent discovery of bacteria within the genus *Nitrospira* capable of complete ammonia oxidation (comammox) demonstrated that the sequential oxidation of ammonia to nitrate via nitrite can also be performed within a single bacterial cell. Although comammox *Nitrospira* exhibit a wide distribution in natural and engineered ecosystems, information on their physiological properties is scarce due to the limited number of cultured representatives. Furthermore, most available genomic information is derived from metagenomic sequencing and high-quality genomes of *Nitrospira* in general are limited. In this study, we obtained a high (90%) enrichment of a novel comammox species, tentatively named “*Candidatus* Nitrospira kreftii”, and performed a detailed genomic and physiological characterization. The complete genome of “*Ca.* N. kreftii” allowed reconstruction of its basic metabolic traits. Similar to *Nitrospira inopinata*, the enrichment culture exhibited a very high ammonia affinity (K_m(app)_NH3_ ≈ 0.036 µM), but a higher nitrite affinity (K_m(app)_NO2_- ≈ 13.8 µM), indicating an adaptation to highly oligotrophic environments. Counterintuitively for a nitrifying microorganism, we also observed an inhibition of ammonia oxidation at ammonium concentrations as low as 25 µM. This substrate inhibition of “*Ca.* N. kreftii” indicate that differences in ammonium tolerance rather than affinity can be a niche determining factor for different comammox *Nitrospira*.

## Introduction

Nitrification, the biological oxidation of ammonia to nitrate via nitrite, is a critical process within the global biogeochemical nitrogen cycle. The nitrification process is of great biotechnological relevance since it fuels the reductive part of the nitrogen cycle and is widely employed in drinking and wastewater treatment systems for the efficient removal of excess ammonium. Traditionally, nitrification was considered to be a two-step process catalyzed by two functionally distinct microbial guilds. According to this paradigm, ammonia-oxidizing prokaryotes first oxidize ammonia to nitrite and subsequently nitrite-oxidizing bacteria (NOB) are responsible for the conversion of nitrite to nitrate. While this dogma has been challenged by the theoretical prediction of complete ammonia oxidation (comammox) (1, 2), it was the discovery of comammox *Nitrospira* that has drastically altered our perception of nitrification (3-5).

All comammox organisms described to date are affiliated with *Nitrospira* sublineage II and can be further divided into clade A and B based on phylogeny of the ammonia monooxygenase, the enzyme catalyzing the first step of ammonia oxidation (4). Comammox *Nitrospira* were identified mainly via metagenomic sequencing in various natural and engineered ecosystems, indicating their widespread occurrence and key role in nitrogen cycling (6-15). This ubiquitous abundance of comammox *Nitrospira* has raised many questions regarding their ecophysiology and potential biotechnological applicability. In order to provide the necessary answers, in-depth understanding of the comammox physiology is required. So far, the sole physiological data available was obtained from *Nitrospira inopinata*, the only existing pure culture of a comammox bacterium (4). The extremely low apparent half saturation constant (K_m(app)_) for ammonia and the high growth yield reported for *N. inopinata* indicate an adaptation to nutrient-limited environments (16) and corroborate the predicted comammox lifestyle (1).

A general adaptation of comammox *Nitrospira* to oligotrophic environments is suggested by their presence mainly in ecosystems with low ammonium loads. However, limited physiological data can highly bias our perception of the ecophysiology of certain microbial groups and kinetic parameters might vary between different comammox species. This was for instance recently observed for ammonia-oxidizing archaea (AOA) and bacteria (AOB), where especially terrestrial AOA were found to have lower ammonia affinities than previously assumed based on the extremely low K_m_ reported for the marine AOA *Nitrosopumilus maritimus* (16, 17). For comammox *Nitrospira*, though, the lack of pure cultures or high enrichments hampers the thorough understanding of the ecophysiology of these intriguing microorganisms.

In this study, we describe the enrichment of a novel comammox *Nitrospira* species in a continuous membrane bioreactor system and provide genome-derived insights into its metabolic potential. Furthermore, we report the ammonia- and nitrite-oxidation kinetics of this comammox organism, including an apparent inhibition by ammonium concentrations as low as 25 µM, findings that provide crucial insights into the niche partitioning factors of different comammox *Nitrospira*

## Materials and Methods

### Enrichment and reactor operation

A 7 L continuous membrane bioreactor (Applikon, Delft, The Netherlands), with a working volume of 5 L was inoculated with biomass (300 mL) from a hypoxic enrichment culture described previously, which contained two distinct comammox *Nitrospira* species (3). The bioreactor was operated for 39 months at room temperature (RT) with moderate stirring (200 rpm). The pH of the culture was constantly monitored by a pH electrode connected to an ADI1020 biocontroller (Applikon, Delft, The Netherlands) and maintained at 7.5 by the automatic supply of a 1M KHCO_3_ solution. Dissolved oxygen was kept at 50% by providing 10 mL/min of a 1:1 mixture of Argon/CO_2_ (95%/5% v/v) and air through a metal tube with a porous sparger. After 450 days of operation a bleed was installed, removing 100 to 300 mL biomass per day.

1 L of sterile NOB mineral salts medium (18) was supplied to the reactor per day. The medium was supplemented with 1 mL of a trace element stock solution composed of NTA (15 g/L), ZnSO_4_·7H_2_O (0.43 g/L), CoCl_2_·6H_2_O (0.24 g/L), MnCl_2_·4H_2_O (0.99 g/L), CuSO_4_·5H_2_O (0.25 g/L), Na_2_MoO_4_·2H_2_O (0.22 g/L), NiCl_2_·6H_2_O (0.19 g/L), NaSeO_4_·10H_2_O (0.021 g/L), H_3_BO_4_ (0.014 g/L), CeCl·6H_2_O (0.24 g/L) and 1 ml of an iron stock solution composed of NTA (10 g/L) and FeSO_4_ (5 g/L). Initially, ammonium, nitrite and nitrate (increasing from 80/0/50 to 250/20/500 µM NH_4_Cl/NaNO_2_/NaNO_3_, respectively) were supplied via the medium. After 60 days of operation, ammonium was supplied as the sole substrate and the concentration was slowly increased to a final concentration of 2.5 mM NH_4_Cl. Liquid samples from the bioreactor were collected regularly for the determination of ammonium, nitrite and nitrate concentrations.

### Analytical methods

Ammonium concentrations were measured colorimetrically via a modified orthophatal-dialdehyde assay (detection limit 10 µM) (19). Nitrite concentrations were determined by the sulfanilamide reaction (detection limit 5 µM) (20). Nitrate was measured by converting it into nitric oxide at 95°C using a saturated solution of VCl_3_ in HCl, which was subsequently measured using a nitric oxide analyzer (detection limit 1 µM; NOA280i, GE Analytical Instruments, Manchester, UK). Protein extraction and determination was performed using the B-PER™ Bacterial Protein Extraction Reagent and Pierce™ BCA Protein Assay Kit (Thermo Fisher Scientific, Waltham, MA, USA), respectively.

### Fluorescence in situ *hybridization*

Biomass samples were fixed using a 3% (v/v) paraformaldehyde (PFA) solution for 30 minutes at RT. Fluorescence *in situ* hybridization (FISH) was performed as described elsewhere (3) using 16S rRNA-targeted oligonucleotide probes (Table S1) that were fluorescently labelled with Fluoresceine or the cyanine dyes Cy3 or Cy5. After hybridization, slides were dried and embedded in Vectashield mounting solution (Vector Laboratories Inc., Burlingame, CA, USA). For image acquisition a Leica TCS SP8x confocal laser scanning microscope (Leica Microsystems, Wetzlar, Germany) equipped with a pulsed white light laser and a 405 nm diode was used. In order to quantify the total *Nitrospira* biovolume in the enrichment culture fixed biomass was hybridized with the probes Ntspa662, Ntspa712 (labelled in the same color) and EUB338mix (Table S1). Subsequently, at least 15 image pairs were recorded at random fields of view. The images were imported into the image analysis software *daime* (21) and analyzed as described elsewhere (22). Similarly, the probes Ntspa1431 and Ntspa1151 in combination with a mix of Ntspa662 and Ntspa712 (Table S1) were used for the biovolume quantification of sublineage I and II *Nitrospira*.

### Floc size determination

A representative biomass sample was used for the determination of the average floc size (area) using image analysis. Microscopic images were acquired using a Zeiss Axioplan 2 (Carl Zeiss AG, Oberkochen, Germany) light microscope. The area of the flocs was calculated manually using the software platform ImageJ (23).

### DNA extraction

After 17 and 39 months of enrichment, DNA was extracted from 50 ml of culture using the PowerSoil DNA Isolation Kit (MO BIO Laboratories Inc., Carlsbad, CA, USA) and a CTAB-based DNA extraction method (17 months sample) (24) or the DNeasy Blood & Tissue Kit (39 months sample; Qiagen, Hilden, Germany). Concentration and quality of the obtained DNA was checked using a Qubit™ dsDNA HS Assay Kit (Thermo Fisher Scientific, Waltham, MA, USA) on a Qubit Fluorometer (Thermo Fisher Scientific, Waltham, MA, USA) and a NanoDrop™ 1000 Spectrophotometer (Thermo Fisher Scientific), respectively.

### Metagenome sequencing and analysis

Metagenome sequencing was performed using an Illumina MiSeq benchtop DNA sequencer (Illumina Inc., San Diego, California USA). Genomic sequencing libraries were prepared using the Nextera XT Kit (Illumina Inc., San Diego, California U.S.A.) following the manufacturer’s instructions, using 1 ng of input DNA normalized to a 0.2 ng/µl concentration. The MiSeq Reagent Kit v.3 (2 x 300 bp) (Illumina Inc., San Diego, California USA) was used for sequencing according to manufacturer’s recommendations.

Sequencing adapter removal, quality-trimming and contaminant filtering of Illumina paired-end sequencing reads was performed using BBDuk version 37.76 from the BBTools package (https://jgi.doe.gov/data-and-tools/bbtools). Processed reads for all samples were co-assembled using metaSPAdes v3.11.1 (25) with default parameters. MetaSPAdes iteratively assembled the metagenome using k-mer lengths 21, 33, 55, 77, 99 and 127. Reads were mapped back to the assembled metagenome for each sample separately using Burrows-Wheeler Aligner (BWA) 0.7.17 (26), employing the “mem” algorithm. The sequence mapping files were processed using SAMtools 1.6 (27). Metagenome binning was performed for contigs ≥2,000 bp. To optimize binning results, five binning algorithms were used: BinSanity v0.2.6.1 (28), COCACOLA (29), CONCOCT 0.4.1 (30), MaxBin 2.0 2.2.4 (31) and MetaBAT 2 2.12.1 (32). To obtain the final bins, the five bin sets subsequently were supplied to DAS Tool 1.0 (33) for consensus binning. The quality of the genome bins was assessed through a single-copy marker gene analysis using CheckM 1.0.7 (34). The GTDB-Tk software was used for taxonomic classifications to the obtained bins (35, 36). Only *Nitrospira* bins with estimated completeness ≥90% and contamination ≤10% were included in subsequent analyses.

### Nanopore sequencing and assembly of “Ca. *N. kreftii”*

To assemble the complete genome of the dominant *Nitrospira* species, single-molecule long-read data was obtained after 17 months of enrichment using the Oxford Nanopore MinION platform (Oxford Nanopore Technologies, Oxford, UK). Genomic DNA was extracted by using the CTAB-based protocol as described above and prepared for sequencing using the Ligation Sequencing Kit 1D (SQK-LSK108, Oxford Nanopore Technologies) according to the manufacturer’s instructions. Adaptor-ligated DNA was cleaned by adding 0.8 volumes of AMPure XP beads (Beckman Coulter Inc., Brea, CA, USA). Sequencing libraries were loaded on a SpotOn Flow Cell (FLO-MIN106 R9.4, Oxford Nanopore Technologies) using the Library Loading Beads Kit (EXP-LLB001, Oxford Nanopore Technologies) following manufacturer’s specifications. Sequencing was performed on the MinION sequencing device with MinKNOW 1.7.10 software using the FLO-MIN106 450 bps protocol. Base calling of the signal data was performed using Guppy v2.3.7 (Oxford Nanopore Technologies) with the flipflop model -c dna_r9.4.1_450bps_flipflop.cfg. Only NanoPore reads with a length ≥700 bp were used for further analyses. The Nanopore reads were assembled *de novo* using Canu v1.8 (37) with parameters “genomeSize=50m” and “corOutCoverage=1000”. Subsequently, all Nanopore reads with a length of ≥700 bp were mapped to the assembly using minimap2 v2.16-r922 (38), followed by building genomic consensus sequences using Racon v1.3.1 (39). This long read assembly approach resulted in a closed *Nitrospira* genome whose taxonomic classification was confirmed using the classify_wf workflow of GTDB-Tk v0.3.2 (40) with default settings. All trimmed Illumina reads of the 17 months sample were mapped to this complete *Nitrospira* genome using bbmap v37.76 (41) with “minid 0.8”. A hybrid assembly was performed using Unicycler v0.4.4 with the mapped Illumina reads and the NanoPore reads as input and the NanoPore consensus assembly as existing long read assembly. In addition, the chromosomal replication initiator protein, DnaA of *N. moscoviensis* (ALA56445.1) was used to set *dnaA* as starting gene with the parameters “start_gene_id 60” and “start_gene_cov 80”.

### Phylogenomic analysis and genome annotation

Reference genomes that were downloaded from the NCBI GenBank database (13/05/2019) and the *Nitrospira* metagenome-assembled genomes (MAGs) retrieved in this study were dereplicated using dRep (33) with default parameters filtering for an estimated completeness ≥90% and contamination ≤10%. The UBCG pipeline (42) was used for phylogenomic analysis of the obtained *Nitrospira* MAGs and 34 publicly available, high-quality genomes of sublineage I and II *Nitrospira*. UBCG was used to identify and extract 91 bacterial single copy core genes from all genomes. After alignment in UBCG with default parameters, a maximum likelihood phylogenetic tree was calculated based on the concatenated nucleotide alignment using RAxML version 8.2.12 (43) on the CIPRES science gateway (44) with the GTR substitution and GAMMA rate heterogeneity models and 100 bootstrap iterations. Two *Leptospirillum* genomes were used as outgroup. Average nucleotide identity (ANI) values of the genomes were calculated using the OrthoANIu tool (45).

All CDS of the high-quality draft genomes of *Nitrospira* including the complete genome of “*Ca.* N. kreftii” were automatically predicted and annotated using a modified version Prokka (46) that performs a BLASTp search of all CDS against the NCBI RefSeq non-redundant protein database (version 92). Homologous proteins in these MAGs and in selected *Nitrospira* genomes were identified by reciprocal best BLAST. Only BLAST hits with an e-value ≤1e-06, amino acid similarity ≥35% and minimum alignment coverage of 80% were considered as homologous proteins. In addition, the automatic annotation of selected genes of *Ca.* N. kreftii was confirmed using the annotation platform Genoscope (47). The visualization tool Circos v0.69-6 (48) was used to generate a whole genome map of “*Ca.* N. kreftii”.

### Adaptation of the enrichment culture in batch culturing conditions with a higher feeding rate

Biomass (1L) from the enrichment culture was washed twice in sterile NOB medium by centrifugation (1500 × *g*, 2 min) and subsequent resuspension in 0.01 M HEPES buffered (pH 7.5) sterile NOB medium containing 0.1 mM KHCO_3_ supplemented with 1 mM NH_4_Cl. The culture was incubated in the dark for 30 days (RT, 150 rpm, rotary shaker). Upon full ammonium consumption, substrate was replenished. Every 15 days the complete culture was centrifuged (1500 × *g*, 2 min) and the medium was exchanged with the same volume of fresh HEPES buffered, sterile NOB medium.

### Substrate-dependent oxygen uptake rate measurements

After 39 months of enrichment, the activity of the culture was determined by microrespirometry. Biomass from 20 mL of the bioreactor or the batch cultures was harvested, washed twice by centrifugation (1500 × *g*, 2 min) and finally resuspended in 2 mL of 0.01 M HEPES buffered sterile NOB medium containing 0.1 mM KHCO_3_. Oxygen consumption rates were measured at 25°C using a RC-350 respiration chamber (Warner Instruments LLC, Hamden, USA), equipped with an oxygen sensor (Model 1302, Warner Instruments LLC, Hamden, USA) and connected to a picoammeter PA2000 (Unisense, Aarhus, Denmark). NH_4_Cl or NaNO_2_ were injected from concentrated stock solutions (1 mM) into the reaction vessel. At the end of the measurements, biomass was harvested for protein determination. Concentrations of ammonium, nitrite and nitrate were determined in the supernatant as described above.

### Calculation of kinetic parameters

The kinetic constants of the enrichment culture were estimated from oxygen consumption measurements using substrate:oxygen consumption stoichiometries of 1:2 and 2:1 for ammonium and nitrite oxidation, respectively. Measurements were corrected for background respiration, which was determined from oxygen uptake rates prior to substrate addition.

Ammonia oxidation by “*Ca.* N. kreftii” was described by a substrate inhibition model and K_i_ values were calculated based on fitting of the data to this model (*Equation 1*). Due to the overestimation of the K_m(app)_ and V_max_ values by the inhibition model, these were obtained by fitting the experimental data obtained for non-inhibitory ammonium concentrations to a Michaelis-Menten model (*Equation 2*), which was also employed to calculate K_m(app)_ and V_max_ for nitrite oxidation.

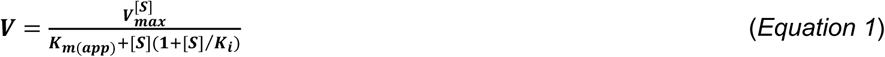

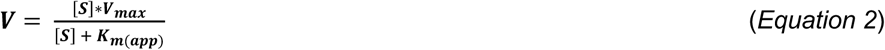

with V representing the observed oxidation rate, V_max_ the maximum rate (in µM h^-1^), K_m(app)_ the apparent Michaelis-Menten half saturation constant, K_i_ the inhibitory half saturation constant and [S] the substrate concentration (in µM).

## Data availability

Sequencing data obtained in this study have been deposited in the National Center for Biotechnology Information (NCBI) database under Bioproject accession number PRJNA575653.

## Results

### Enrichment of comammox Nitrospira

A continuous laboratory-scale membrane bioreactor was used for the enrichment of comammox *Nitrospira*. The bioreactor was inoculated with biomass from an enrichment culture containing two comammox *Nitrospira* species (“*Ca.* Nitrospira nitrosa” and “*Ca.* Nitrospira nitrificans”) that constituted together approximately 15% of the microbial community (3). Since comammox bacteria are speculated to thrive under highly limiting substrate concentrations, medium amended with low ammonium concentrations was supplied to the system. The total ammonium load of the system was, based on the consumption rate and biomass concentration in the culture, gradually increased from initially 0.016 to 2.5 mmol/day (Figure S1A) and was stoichiometrically oxidized to nitrate (Figure S1B). Concentrations of ammonium and nitrite in the bioreactor always remained below the detection limit (10 µM; Figure S1B).

After 27 months of operation, *Nitrospira* bacteria were present in suspended flocs of an average area of 9.8 µm^2^ and constituted approximately 90% of the total microbial community in the culture, as revealed by quantitative FISH (Figure 1, Table S2). Subsequently, the relative abundance of *Nitrospira* dropped, but they remained the dominant community member over the whole 39 months of operation.

**Figure 1.**
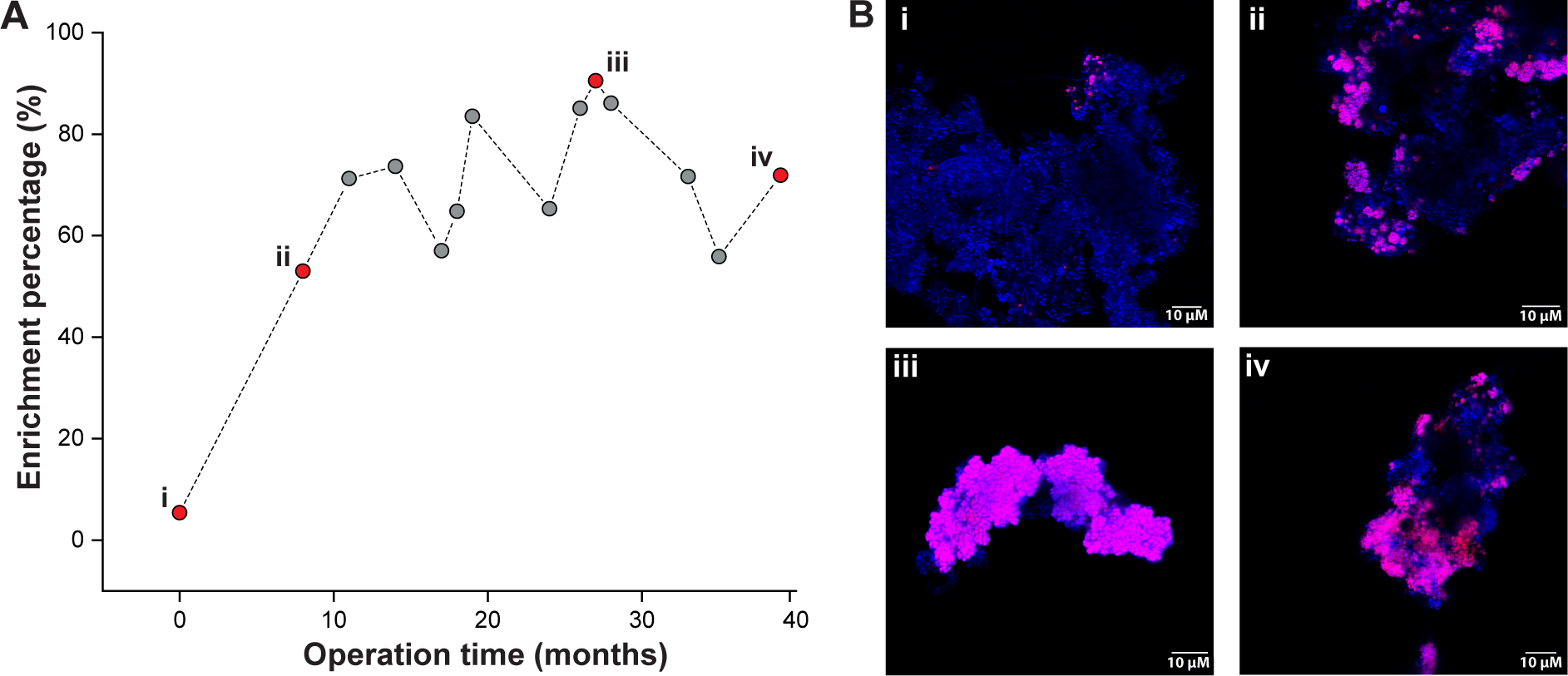
Enrichment of *Nitrospira* bacteria in the bioreactor system. (A) Relative abundance of *Nitrospira* bacteria over the enrichment period. (B) Representative fluorescent micrographs of biomass samples obtained from the enrichment culture throughout the enrichment period. i) Starting inoculum of the bioreactor and biomass sampled after ii) 8, iii) 26 and iv) 39 months of enrichment. Cells are stained using FISH probes for all bacteria (EUB338mix, blue) and *Nitrospira*-specific probes (Ntspa712 and Ntspa662, red).

Quantification of the relative abundances of *Nitrospira* affiliated with sublineages I and II revealed that up to 95 ± 6% of the total *Nitrospira* population consisted of sublineage II *Nitrospira*, while sublineage I never constituted more than 3.1 ± 1% (Figure S2). FISH with probes targeting the known AOA or betaproteobacterial AOB indicated their absence from the culture at all time points analyzed (data not shown), as was already the case for the inoculum (3).

### Metagenomic retrieval of a novel clade A comammox Nitrospira

Metagenome sequencing in combination with *de novo* assembly and consensus binning of the microbial community present in the bioreactor enrichment after 17 months of operation resulted in the recovery of 28 metagenome-assembled genomes (MAGs) of medium or high quality (completeness ≥75% or ≥90%, respectively, and contamination ≤10%; Dataset S1). Of these, four high-quality MAGs were affiliated with the genus *Nitrospira*. The number of reads mapped to these *Nitrospira* MAGs corresponded to 36% of the total reads and total coverage data suggested that one *Nitrospira* MAG dominated the microbial community in the bioreactor system (Dataset S1). Phylogenomic analysis revealed that this MAG belongs to a novel clade A comammox *Nitrospira* (Figures 2 and S3). The remaining *Nitrospira*-like MAGs clustered with canonical nitrite-oxidizing *Nitrospira* within sublineage I (2 MAGs; *Nitrospira* spp. CR1.1 and CR1.2) and sublineage II (1 MAG; *Nitrospira* sp. CR1.3; Figures 2 and S3). In combination with the lack of key genes for ammonia oxidation (Figure S4), this phylogenetic affiliation strongly indicated that these *Nitrospira* were canonical nitrite oxidizers.

**Figure 2.**
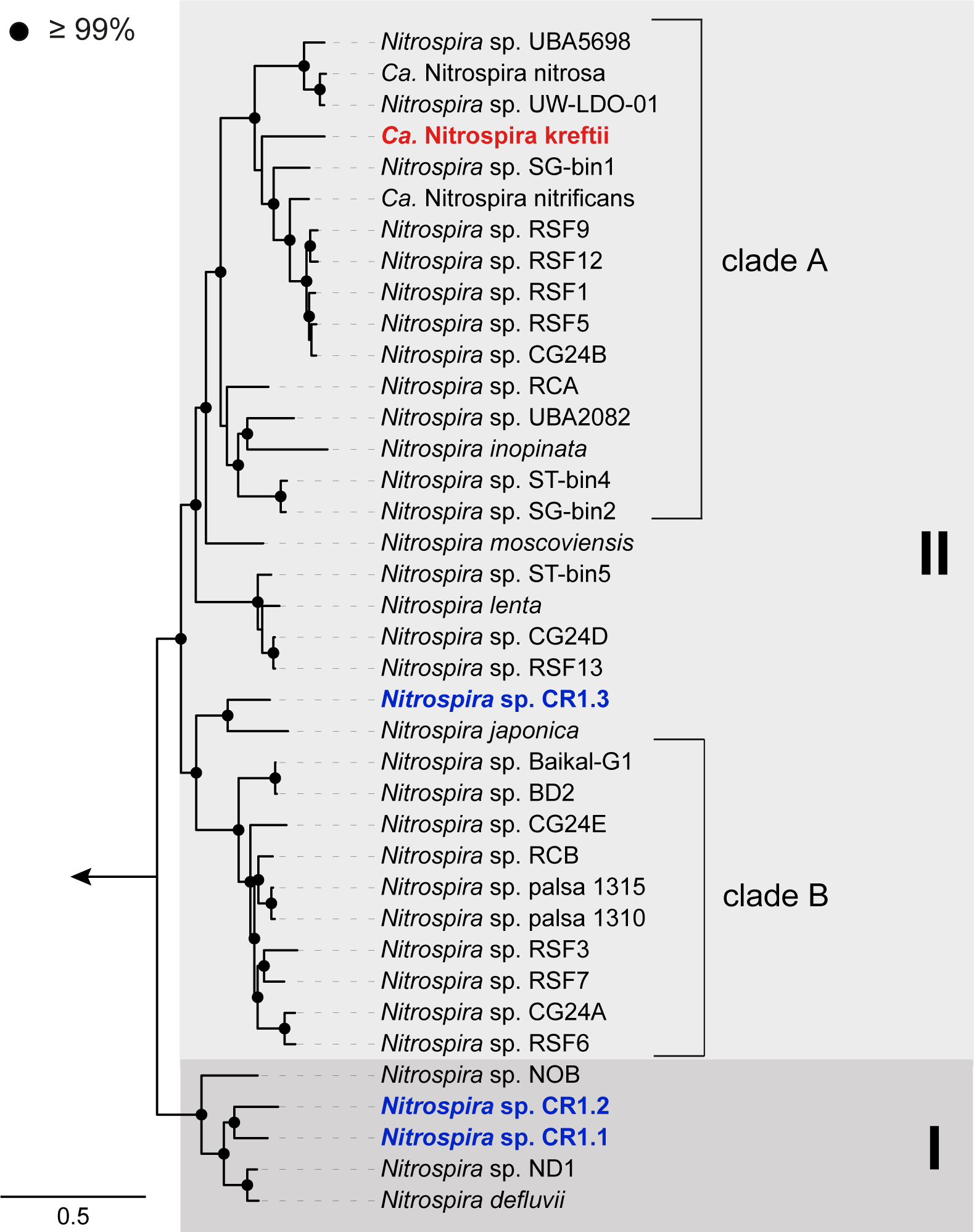
Phylogenomic analysis of the retrieved *Nitrospira* MAGs and representative sublineages I and II *Nitrospira*. Sublineages are indicated by shaded boxes and labeled with roman numerals, comammox clades are designated by square brackets. Bootstrap support values ≥99% are indicated by black circles. The arrow indicates the position of the outgroup, which consisted of two *Leptospirillum* species. The scale bar corresponds to 50% estimated sequence divergence.

A hybrid assembly approach for scaffolding the Illumina assembly with long Nanopore reads allowed the retrieval of the complete genome of the dominant *Nitrospira* MAG, with a total size of 4.13 Gb and an overall G+C content of 54.5%. The average nucleotide identities (ANI) of this genome to 34 high-quality genomes of sublineage I and II *Nitrospira* available at the time of study (May 13, 2019) are ≤77% (Figure S3), which is below the species cut off of 95% (49). Together with the phylogenetic distance in the phylogenomic analysis (Figure 2), this classifies it as a novel clade A comammox *Nitrospira*, which we tentatively named ‘*Candidatus* Nitrospira kreftii’.

Resequencing after additional 22 months of enrichment indicated a clear decrease in diversity for both *Nitrospira* and the overall microbial community. More specifically, after a total of 39 months of enrichment, we retrieved 9 medium and 7 high-quality MAGs, out of which “*Ca.* N. kreftii” was the only MAG affiliated with sublineage II of the genus *Nitrospira* (Dataset S2). The metagenome contained one additional MAG (*Nitrospira* sp. CR2.1) representing a canonical sublineage I *Nitrospira*, which was highly similar (96% ANI) to the *Nitrospira* sp. CR1.2 MAG obtained from the 17-month sample. However, this MAG showed >10% estimated contamination, most likely indicating wrong assignments of contigs belonging to *Nitrospira* sp. CR1.1 into this genome bin. Putatively heterotrophic microorganisms accounted for the rest of the microbial community present in the system (Dataset S2). In both metagenomic datasets no canonical ammonia-oxidizing prokaryotes were identified, confirming that “*Ca.* N. kreftii” was indeed the only ammonia oxidizer in the system (Datasets S1 and S2).

### Metabolic potential of “Ca. *N. kreftii”*

Analysis of the complete “*Ca.* N. kreftii” genome revealed the presence of all genes for the enzyme complexes involved in complete nitrification (i.e., ammonia monooxygenase (AMO), hydroxylamine dehydrogenase (HAO) and nitrite oxidoreductase (NXR); Figure 3, Dataset S3). Similar to most comammox *Nitrospira*, the *Ca.* N. kreftii genome contained one gene cluster encoding the structural AMO subunits (*amoCAB*), the hydroylamine:ubiquinone reduction module (HURM; consisting of *haoAB* for the HAO structural subunits and *cycAB*, encoding the cytochromes *c*554 and *c*_M_552) as well as genes for the type I cytochrome *c* biosynthesis system. In addition, “*Ca.* N. kreftii” harbors four non-operonal *amoC* copies and an additional *haoA* (Dataset S3). For nitrite oxidation, the genome contains two *nxrAB* gene clusters encoding the alpha and beta subunit of the periplasmic NXR and four non-operonal genes for putative gamma subunits (*nxrC*; Figure S5). One of these *nxrC* is clustered with a TorD-like chaperone probably involved in insertion of the molybdopterin cofactor into the catalytic NxrA subunits, and a NapG-like ferredoxin as has been described for other *Nitrospira* (50-52). As found in all *Nitrospira* (53, 54), also “*Ca.* N. kreftii” encodes a copper-containing nitrite reductase (NirK; Dataset S3), the exact function of which however still is unclear (16).

**Figure 3.**
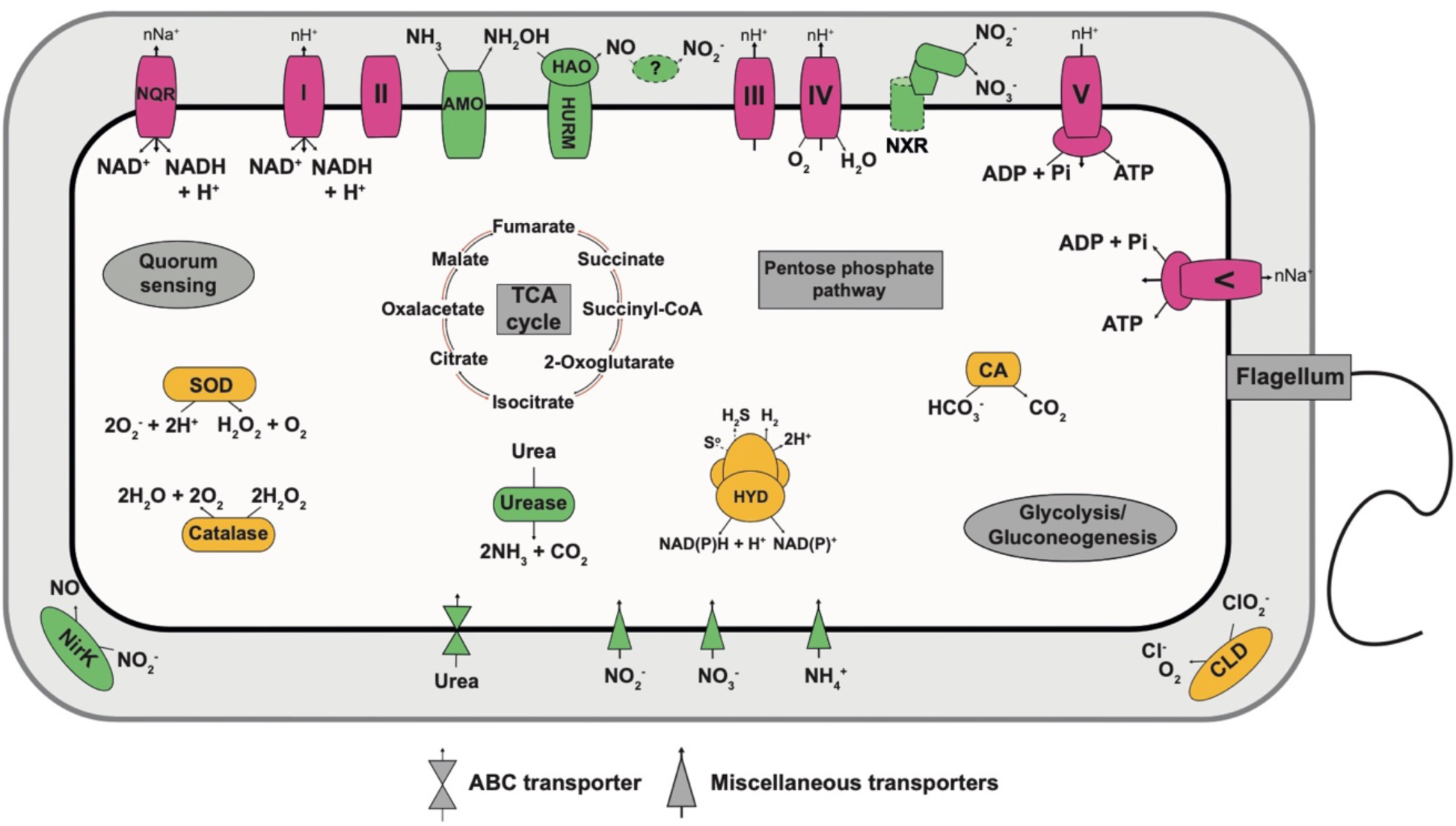
Cell metabolic cartoon of “*Ca.* N. kreftii”. AMO: ammonia monooxygenase, HAO: hydroxylamine dehydrogenase, NXR: nitrite oxidoreductase, HYD: group 3b [NiFe]-hydrogenase, CA: carbonic anhydrase, CLD: chlorite dismutase, SOD: superoxide dismutase, NirK: Cu-dependent nitrite reductase, NQR: Na^+^-translocating NADH:ubiquinone oxidoreductase. Enzyme complexes of the electron transport chain are labeled by Roman numerals. Dashed lines indicate putative features. The question mark indicates that the exact enzyme catalyzing the nitrite formation from NO remains uncertain.

In addition to the enzyme systems for ammonia and nitrite oxidation, “*Ca.* N. kreftii” encodes all complexes of the respiratory chain, the reductive tricarboxylic acid cycle for CO_2_ fixation, and the complete gene repertoire for glycolysis/gluconeogenesis and the oxidative and non-oxidative phases of the pentose phosphate pathway, which all belong to the core metabolism of the genus *Nitrospira*. Notably, the genome of “*Ca.* N. kreftii” encoded an alternate F_1_F_o_-type ATPase (also referred to as Na^+^-translocating N-ATPase; (55)), and an alternative sodium-pumping complex I (Na^+^-NQR; Dataset S3) (56). These features have been identified in other aerobic and anaerobic ammonia oxidizers, as well as in the nitrite-oxidizing “*Ca.* Nitrospira alkalitolerans”, where they were linked to an adaptation to haloalkaline environments (57-60). Similar to other clade A comammox *Nitrospira* (3, 16, 61), “*Ca.* N. kreftii” featured a low-affinity Rh-type transporter for ammonium uptake and possessed a complete operon encoding the structural and accessory urease subunits, and a high-affinity urea transporter. Besides urea, canonical *Nitrospira* use nitrite as nitrogen source for assimilation (Figure S4).

While canonical *Nitrospira* can use nitrite as nitrogen source for assimilation (Figure S4), no assimilatory nitrite reduction system was identified in the complete genome of “*Ca.* N. kreftii” (Dataset S3), as is the case in all other available comammox genomes (13).

### Kinetic characterization of the enrichment culture

Both FISH and metagenomic sequencing indicated the absence of known canonical ammonia oxidizers in the enrichment culture, and demonstrated “*Ca.* N. kreftii” to be the dominant nitrifier and only comammox *Nitrospira* in the system. Thus, the enrichment culture was used to determine the apparent kinetic parameters of the nitrification reaction by measuring the substrate-dependent oxygen (O_2_) uptake rates using microrespirometry.

O_2_ consumption immediately increased upon substrate addition and ammonium and O_2_ were consumed in a 1:2 stoichiometry (1:1.96 ± 0.13 mean ± s.d., n=4), as expected for complete nitrification. From this data we estimated an apparent half-saturation constant (K_m(app)_) of 2.03 ± 0.56 µM total ammonium (NH_3_ + NH_4_^+^; n=3), corresponding to K_m(app)_ ≈ 0.036 ± 0.01 µ ammonia. The maximum total ammonium oxidation rate (V_max_) of 85.8 µmol ± 15.2 N (mg protein)^-1^ hour^-1^ (n=3) was reached at ammonium concentrations as low as 25 µM (Figures 4 and S6). Surprisingly however, ammonia oxidation by the enrichment culture did not follow typical Michaelis–Menten kinetics and ammonium concentrations >25 µM caused a reduction in V_max_. Consequently, ammonia oxidation *in* “*Ca*. N. kreftii” was better described using a substrate inhibition model (see Materials and Methods), which yielded an apparent inhibition constant (K_i(app)_) of 261.6 ± 98.7 µM total ammonium (n = 3; Figures 4 and S6), corresponding to K_i(app)_ ≈ 4.65 ± 1.8 µM ammonia.

**Figure 4.**
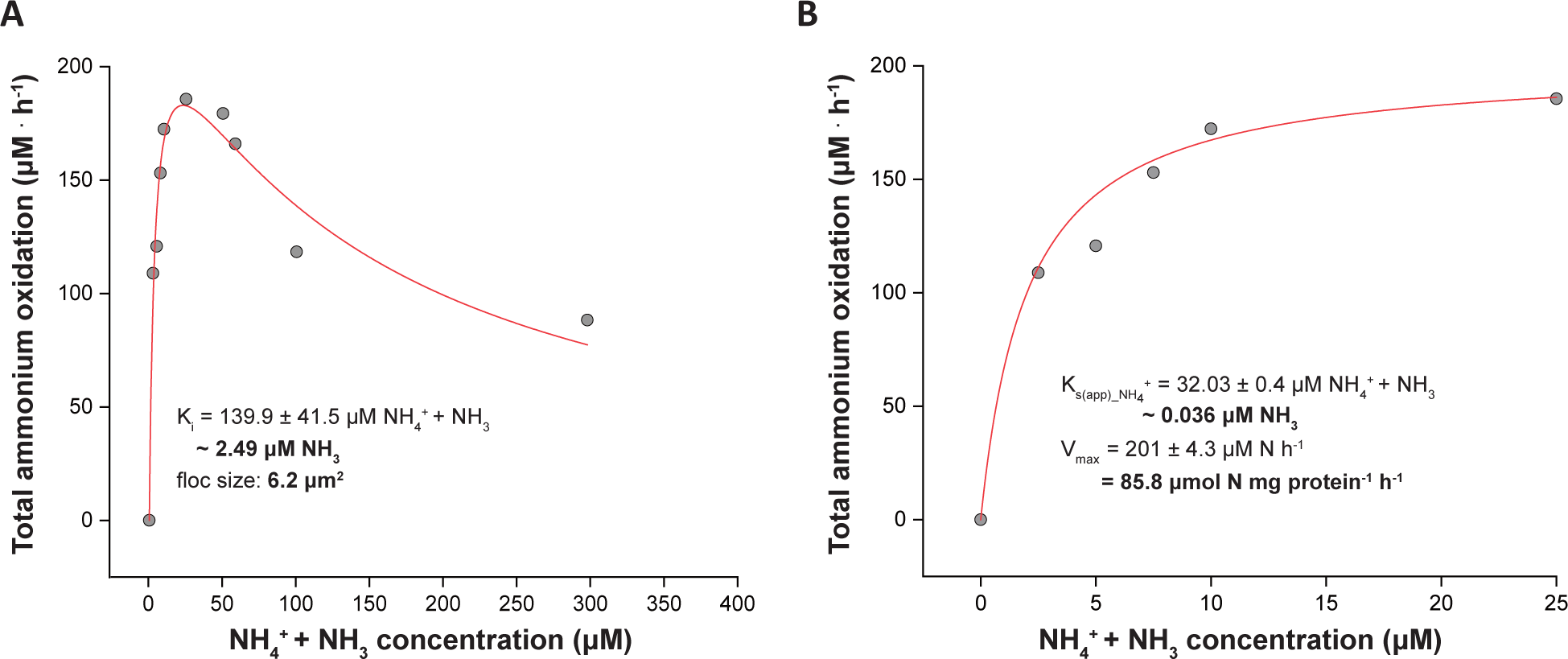
Ammonia oxidation kinetics of the “*Ca.* N. kreftii” enrichment culture. The red curves indicate the best fit (A) of all data to the substrate inhibition model and (B) of the data retrieved for non-inhibitory ammonium concentrations in a Michaelis-Menten kinetic equation. The reported standard errors are based on nonlinear regression. Data from additional biological replicates are shown in Figure S6.

Contrastingly, nitrite oxidation in the enrichment culture followed typical Michaelis-Menten kinetics. Nitrite was oxidized to nitrate with the expected 2:1 nitrite:oxygen stoichiometry (2:1.04 ± 0.04, n=3) a K_m(app)_ = 13.8 ± 4 µM nitrite (n = 3) and V_max_ = 57.8 ± 2.1 µM nitrite (mg protein)^-1^ hour^-1^ (n=3) were estimated (Figures 5 and S7). For non-planktonic microbial cultures, substrate uptake kinetics are influenced by the size and shape of the microcolonies the microorganism forms, and the thickness of the biofilm or, in case of suspended growth, floc size (62). The average floc size of the biomass during determination of the ammonia and nitrite oxidation kinetic parameters was 4.9 µm^2^ (1.6 – 8.4 µm^2^). As expected, the observed K_m(app)_ values varied with floc size, with increasing floc sizes corresponding to larger K_m(app)_ values (Figures 4, 5, S6 and S7).

**Figure 5.**
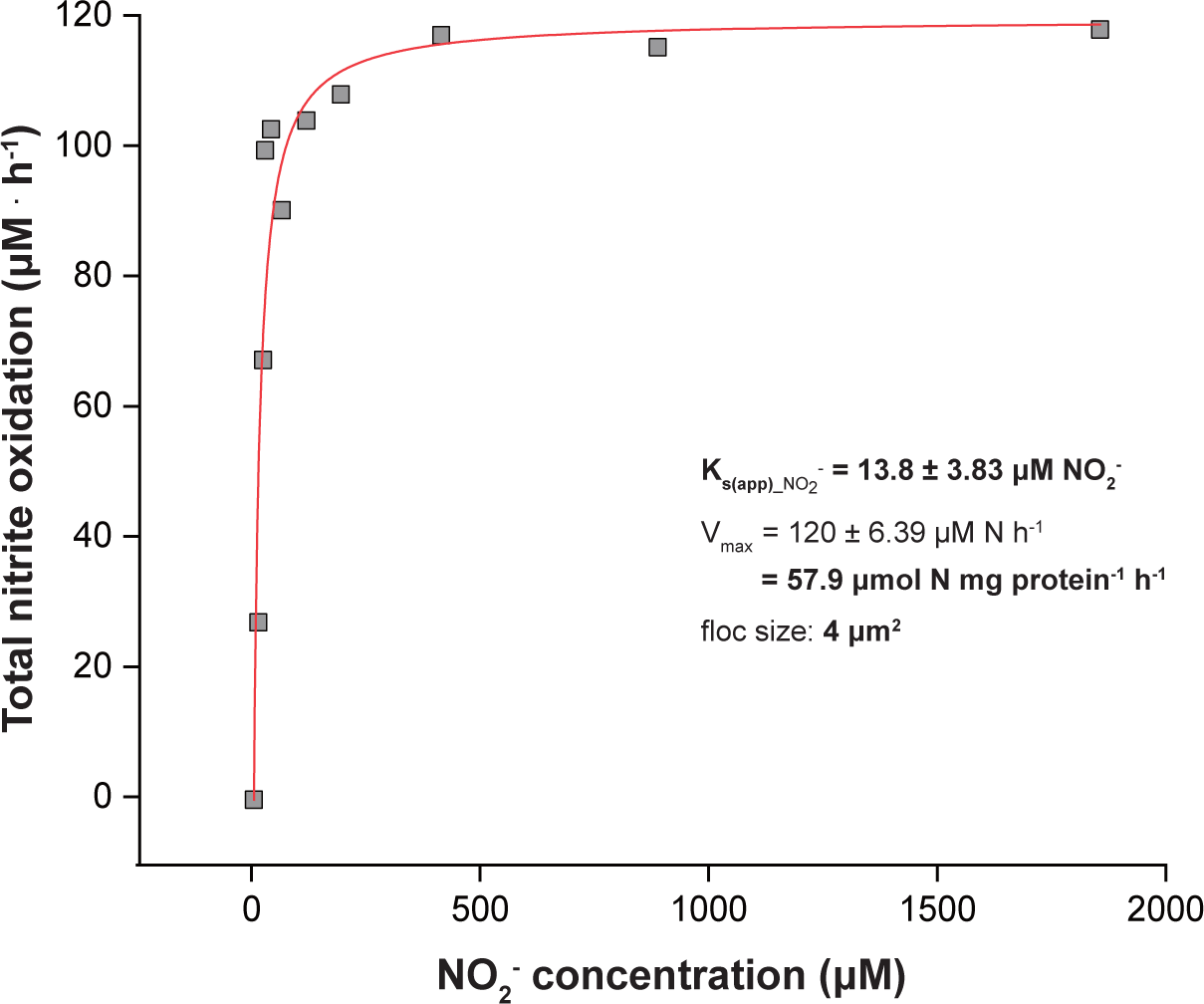
Nitrite oxidation kinetics of the “*Ca.* N. kreftii” enrichment culture. The red curve indicates the best fit of the data to the Michaelis-Menten kinetic equation. The reported standard errors are based on nonlinear regression. Data from additional biological replicates are shown in Figure S7.

To exclude that the observed inhibition pattern of ammonia oxidation was due to a potential physiological adaptation of the biomass to the continuous substrate-limited culturing conditions, batch cultures at higher ammonium concentrations (1 mM NH_4_^+^) were initiated. After 30 days of cultivation with substrate replenishment whenever ammonium was fully consumed, ammonia and nitrite oxidation kinetics were determined as before. However, also with this high substrate-adapted biomass, a highly similar inhibition pattern was observed upon addition of ammonium concentrations >25 µM (Figures 6 and S8), whereas the nitrite oxidation kinetics again followed typical Michaelis-Menten type kinetics (Figures 6 and S9). Fitting of the converted oxygen uptake data to *Equations 1* and *2* (see Materials and Methods) yielded K_m(app)_, V_max_ and K_i(app)_ values for ammonia and nitrite oxidation that where almost identical to those obtained with the continuous bioreactor culture.

**Figure 6.**
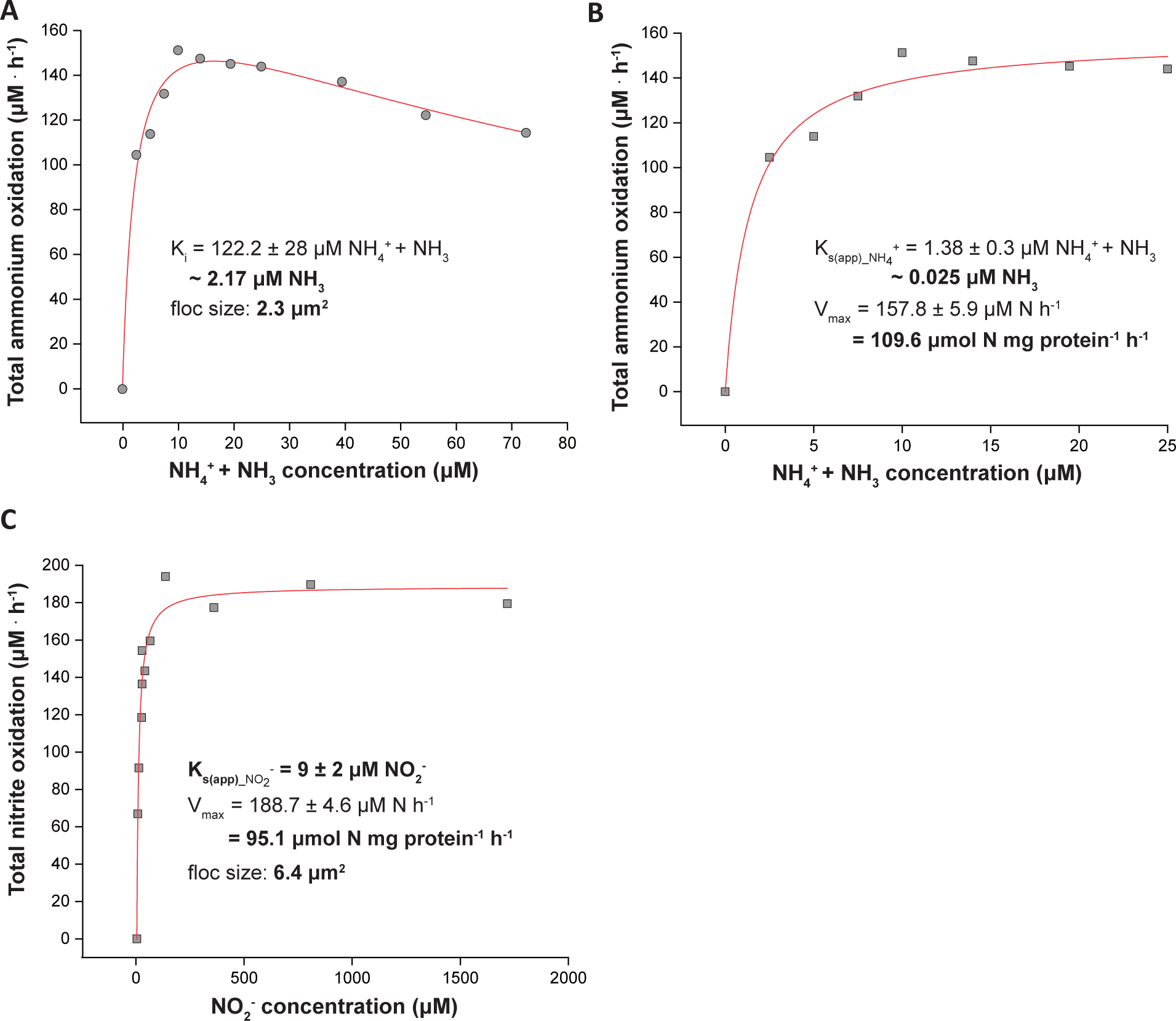
Substrate oxidation kinetics of the “*Ca.* N. kreftii” enrichment culture adapted to 1mM ammonium feeding. (A, B) Ammonia oxidation kinetics; the red curves indicate the best fit (A) of all data to the substrate inhibition model and (B) of the data retrieved for non-inhibitory ammonium concentrations in a Michaelis-Menten kinetic equation. Data from additional biological replicates are shown in Figure S8. (C) Nitrite oxidation kinetics; the red curve indicates the best fit to the Michaelis-Menten kinetic equation. All the reported standard errors are based on nonlinear regression. Data from additional biological replicates are shown in Figure S9.

Lastly, the inhibitory effect of elevated ammonium concentrations on “*Ca*. N. kreftii” was verified in batch incubations. All incubations were inoculated with biomass from the bioreactor system and amended with different amounts of substrate. Addition of >100 µM ammonium caused a lag phase until ammonia oxidation was initiated (Figure S10A), and maximum activity was observed to also decrease (Figure S10B). Contrastingly, nitrite oxidation rates continued to increase at higher nitrite concentrations in batch incubations (Figure S10D) and no effect of the initial nitrite concentration on the lag phase of the culture was observed (Figure S10C).

## Discussion

Recent metagenomic studies have demonstrated the abundance of comammox *Nitrospira* in numerous natural and engineered ecosystems, hinting at their crucial role within the biogeochemical nitrogen cycle (12). However, the ecological niche of these enigmatic organisms is still unclear. Theoretical kinetic modeling studies have predicted that comammox organisms would thrive in environments that select for low growth rates and high yields, as for instance encountered in biofilm-like systems under substrate-limited conditions (1, 2). First physiological data of the pure culture *N. inopinata* substantiated these predictions, as this comammox *Nitrospira* was shown to have an extremely high ammonia affinity and growth yield, which even surpassed those of terrestrial AOA (16). This indicates that in highly oligotrophic environments not AOA but comammox *Nitrospira* could be the main drivers of nitrification. However, the limited availability of cultured representatives still hinders full appreciation of the unique comammox ecophysiology and thus additional cultures are urgently needed in order to fully understand their contribution to nitrification in the environment and their potential biotechnological applicability.

Here, a novel comammox *Nitrospira* species was highly enriched in a continuous substrate-limited bioreactor system. The enrichment culture performed complete nitrification without transient nitrite accumulation (Figure S1). Metagenomic sequencing after 17 months of bioreactor operation revealed the presence of three canonical nitrite-oxidizing and one clade A comammox *Nitrospira* species (Figure 2). Additional long-read sequencing facilitated the reconstruction of the complete genome of this comammox *Nitrospira*, which, based on pairwise ANI comparisons (Figure S3) and phylogenetic distance (Figure 2), forms a novel species tentatively named “*Ca.* N. kreftii”. Notably, this genome represents only the second complete genome available for comammox *Nitrospira*.

Genome analysis indicated high metabolic overlap with the phylogenetically closely related “*Ca.* N. nitrificans”, with which it also shares the highest genome identity (77% ANI). Like all comammox *Nitrospira*, they share the enzymatic machineries required for energy conservation by ammonia and nitrite oxidation. While the complexes of the respiratory chain including the periplasmic NXR are conserved among all *Nitrospira* (13), the key enzymes for ammonia and hydroxylamine oxidation are confined to comammox *Nitrospira* and have highest similarity to the respective enzymes in betaproteobacterial AOB (3, 4, 54). For these it has recently been proposed that nitric oxide (NO) is an obligate intermediate of the ammonia oxidation process (63). In this revised model, NO is produced by HAO and subsequently oxidized to nitrite abiotically or, more likely, enzymatically. One of the best candidates for NO oxidation is the NO-forming nitrite reductase NirK, which would operate in reverse during aerobic ammonia oxidation (63). NirK is conserved in all *Nitrospira* including “*Ca.* N. kreftii”, but its function in the ammonia oxidation pathway remains to be verified.

The co-occurrence of canonical nitrite-oxidizing and comammox *Nitrospira* in the enrichment culture (Figures 2 and S2) indicates a functional relationship between the two microorganisms in the system. Despite the fact that nitrite remained always below the detection limit (<5 µM) in the bioreactor, previous studies on *N. inopinata* have shown the transient accumulation of nitrite in comammox batch cultures (16). Thus, comammox *Nitrospira* might always excrete some nitrite during ammonia oxidation, which in mixed culture systems might immediately be consumed by canonical *Nitrospira* with higher nitrite affinities (16). This would explain the presence of canonical *Nitrospira* in the enrichment and indicate an unexpected potential interplay between the two functional types of *Nitrospira* similar to the symbiotic interactions between canonical AOB and NOB (64), with nitrite-oxidizing *Nitrospira* relying on leakage of nitrite from comammox *Nitrospira*.

Besides canonical *Nitrospira*, metagenomic sequencing furthermore indicated the perseverance of a complex microbial community, consisting mainly of potential heterotrophic microorganisms. Thus, despite the high degree of enrichment of “*Ca.* N. kreftii” achieved, a combination of physical separation and traditional microbiological techniques appears necessary to obtain a pure culture from the bioreactor’s biomass. Several protocols, including label free cell sorting (65, 66), optical tweezers (67) and very recently an automated Raman-based microfluidics platform (68) could assist in the future isolation of “*Ca.* Nitrospira kreftii”. However, while pure cultures are of undoubtful importance for a thorough physiological characterization of an organism, also enrichment cultures can provide invaluable insights into their ecophysiology. When we investigated the ammonia oxidation kinetics of our “*Ca.* N. kreftii” enrichment, we determined a very high ammonia affinity (K_m(app)_NH3_ ≈ 0.036 ± 0.01 µM, n=3). However, this value must be considered as a conservative approximation, as diffusion limitations due to the flocculation of the biomass (average floc size 5.9 µm^2^) are expected to have caused a strong underestimation of the substrate affinity. Correspondingly, when repeated with less aggregated biomass (average floc size 3 µm^2^), a lower substrate affinity was measured (K_m(app)_NH3_ ≈ 0.032 µM), and the opposite was observed in experiments with larger flocs (average floc size 8.4 µm^2^, K_m(app)_NH3_ ≈ 0.051 µM; Figure S6). These values are very similar to the reported ammonia affinity of *N. inopinata* (16) and confirm that comammox *Nitrospira* exhibit a substrate affinity orders of magnitude higher than most characterized AOB and even one order higher than non-marine AOA (Figure 7A). The high ammonia affinity determined for “*Ca.* N. kreftii” agrees well with previous theoretical predictions of the comammox ecophysiology (1, 2) and further verifies an adaptation of comammox bacteria to extremely oligotrophic environments (16).

**Figure 7.**
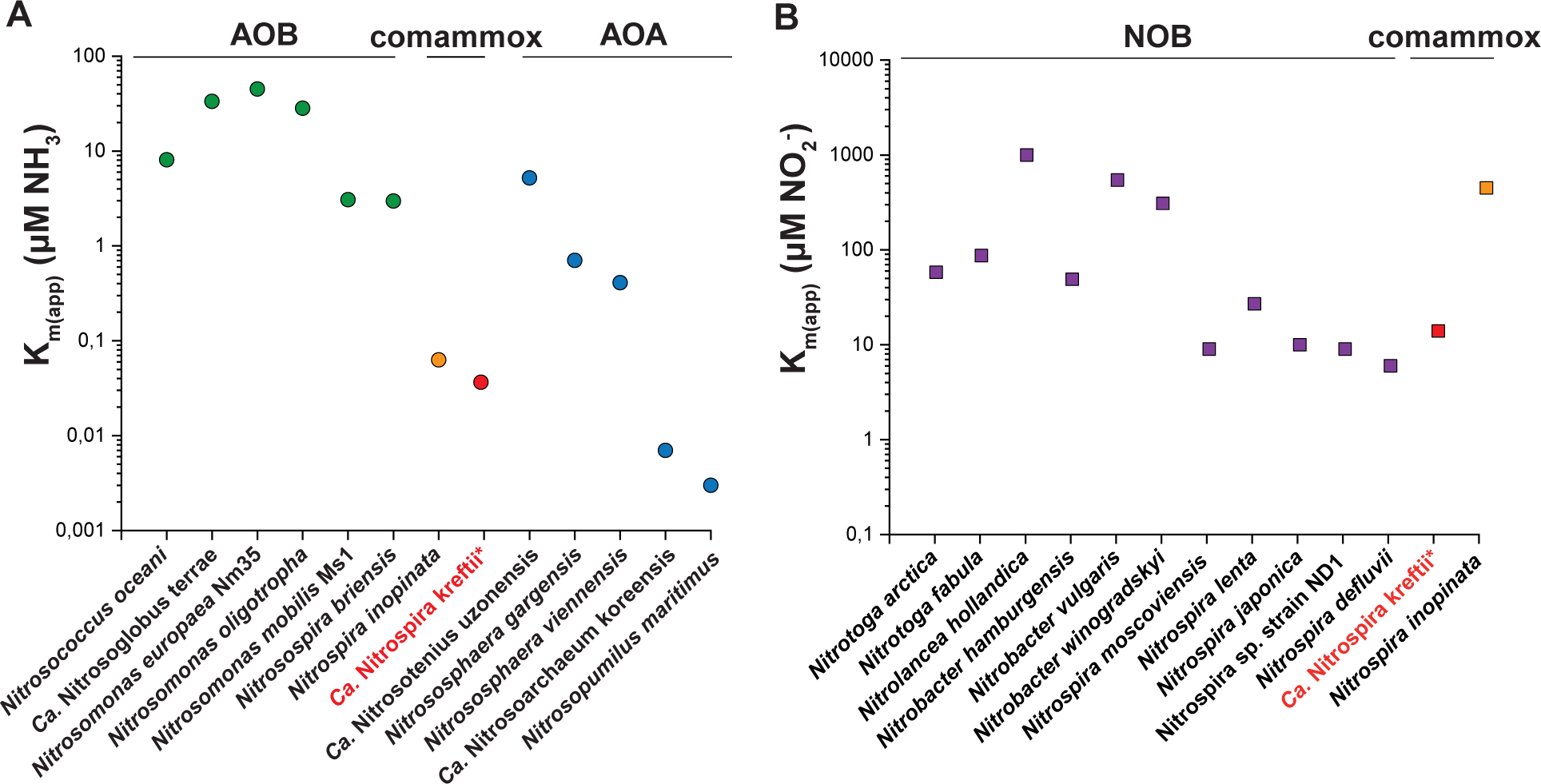
Apparent affinity constants for (A) ammonia and (B) nitrite of the “*Ca.* N. kreftii” enrichment culture (red symbols) in comparison to the reported values of *N. inopinata* (orange) and canonical AOA (blue), AOB (green) and NOB (purple) (16, 17, 38, 71-79). When ammonia affinity values were not given in the respective studies, these were calculated from the reported total ammonium concentrations, pH and temperature provided. The asterisk indicates that the highly enriched “*Ca.* N. kreftii” culture contains also canonical, nitrite-oxidizing *Nitrospira*.

Surprisingly, already very low ammonium concentrations (>25 µM) were found to have an inhibitory effect on ammonia oxidation by the “*Ca.* N. kreftii” enrichment. Although a similar ammonium inhibition pattern was not observed for *N. inopinata* (16), ammonium-sensitive canonical AOB affiliated with *Nitrosomonas* group 6a have been isolated previously (69, 70) and ammonia-induced inhibition of nitrifying microorganisms in activated sludge and soil has been described as well (71, 72). This inhibition of some ammonia-oxidizing microorganisms is thought to be a consequence of their adaptation to substrate-limited environments, or alternatively as sensitivity to the toxic effects of free ammonium itself or to intermediates of the ammonia oxidation pathway (70, 72). However, it was not possible to adapt the “*Ca.* N. kreftii” enrichment culture, as pre-incubation at higher ammonium concentrations (1 mM) did not result in a reduction of the inhibitory effects (Figure 6). Moreover, in batch incubations with biomass from the enrichment culture, a prolonged lag phase was observed when incubated in the presence of >100 µM ammonium (Figure S10), indicating that this adaptation of “*Ca.* N. kreftii” to extremely low substrate concentrations was independent of the method used to study its ammonia oxidation kinetics and could not be attributed to continuous culturing under substrate-limited conditions.

Nitrite oxidation in the “*Ca.* N. kreftii” enrichment followed canonical Michaelis-Menten kinetics and a substrate affinity consistent with canonical nitrite-oxidizing *Nitrospira* was obtained (K_m(app)_NO2_- = 13.7 ± 4 µM, n=3, average floc size 3.9 µm^2^, Figure 5). As this value was determined in a system containing comammox and canonical nitrite-oxidizing *Nitrospira*, this represents the combined affinity of the two functionally distinct *Nitrospira* types. However, the low relative abundance of canonical nitrite-oxidizing *Nitrospira* at the time these experiments were conducted (3.1% of the total *Nitrospira* population; Figure S2) suggests that also “*Ca*. N. kreftii” exhibits this high nitrite affinity, which is in stark contrast to *N. inopinata* (K_m(app)_ NO2_-= 449.2 ± 65.8 µM) (16). These physiological differences between comammox species emphasize the need for the investigation of several representatives of a microbial guild in order to obtain a complete picture of its ecophysiological potential.

In conclusion, the obtained enrichment culture enabled the genomic and physiological characterization of the novel comammox species “*Ca.* N. kreftii”. While there were only few metabolic differences predicted by genomic analyses compared to other clade A comammox *Nitrospira*, clear deviations were observed to *N. inopinata* regarding their ammonia and nitrite oxidation kinetics. The apparently higher substrate affinities of “*Ca.* N. kreftii” for ammonia and nitrite compared to canonical AOB, non-marine AOA, and *N. inopinata*, respectively, indicated a physiological advantage in highly oligotrophic environments. Furthermore, the observed inhibition by ammonium indicated differences in substrate tolerance of comammox *Nitrospira* that could play a crucial role in their interspecies competition and ecological niche partitioning. These novel insights into the physiology of comammox *Nitrospira* further expand our understanding of these enigmatic microorganisms and can have great implications on process designing for their biotechnological application.

### Taxonomic consideration of ‘*Candidatus* Nitrospira kreftii’ *sp. nov.*

N.L. gen. n. kreftii, of Kreft, in honor of Jan-Ulrich Kreft, a German computational biologist, for his leading contribution to the theoretical prediction of comammox bacteria. Phylogenetically affiliated with sublineage II of the genus *Nitrospira*. Belongs to comammox clade A; capable of complete nitrification.

## Acknowledgments

We would like to thank Theo van Alen, Laura Wenzel and Guylaine Nuijten for technical assistance. The LABGeM (CEA/IG/Genoscope & CNRS UMR8030) and the France Génomique National infrastructure (funded as part of Investissement d’avenir program managed by Agence Nationale pour la Recherche, contract ANR-10-INBS-09) are acknowledged for support within the MicroScope annotation platform. We are grateful to Lianna Poghosyan, Arjan Pol and Huub Op de Camp for helpful discussions. D.S. and M.S.M.J. were supported by the European Research Council (ERC Advanced Grant Ecomom 339880), J.F., M.A.H.J.v.K and S.L. by the Netherlands Organisation for Scientific Research SIAM (NWO; Gravitation Grant SIAM 024.002.002, 016.Veni.192.062 and 016.Vidi.189.050, respectively). H.K. gratefully acknowledges the Radboud Excellence Initiative.

## Author Contributions

S.L. conceived the presented research. D.S., M.A.H.J.v.K, M.S.M.J. and S.L. planned the research. S.L. and M.A.H.J.v.K supervised the project. D.S., H.K. and J.F. executed experiments and analyzed data. D.S., M.A.H.J.v.K and S.L. wrote the paper. All authors discussed results and commented on the manuscript.

