## Supplemental Material for "Enrichment and physiological characterization of a novel comammox *Nitrospira* indicates ammonium inhibition of complete nitrification"

### Supplemental figures and tables

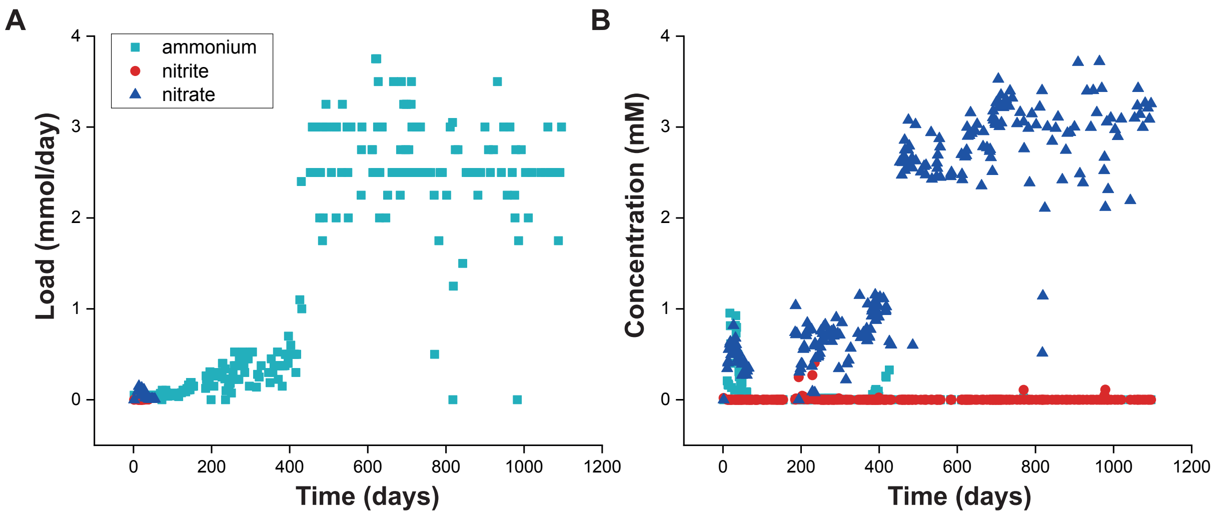


**Figure S1.** Substrate loading (A) and apparent substrate concentrations (B) in the bioreactor system during the enrichment period. Symbols indicate ammonium (squares), nitrite (circles) and nitrate (triangles).

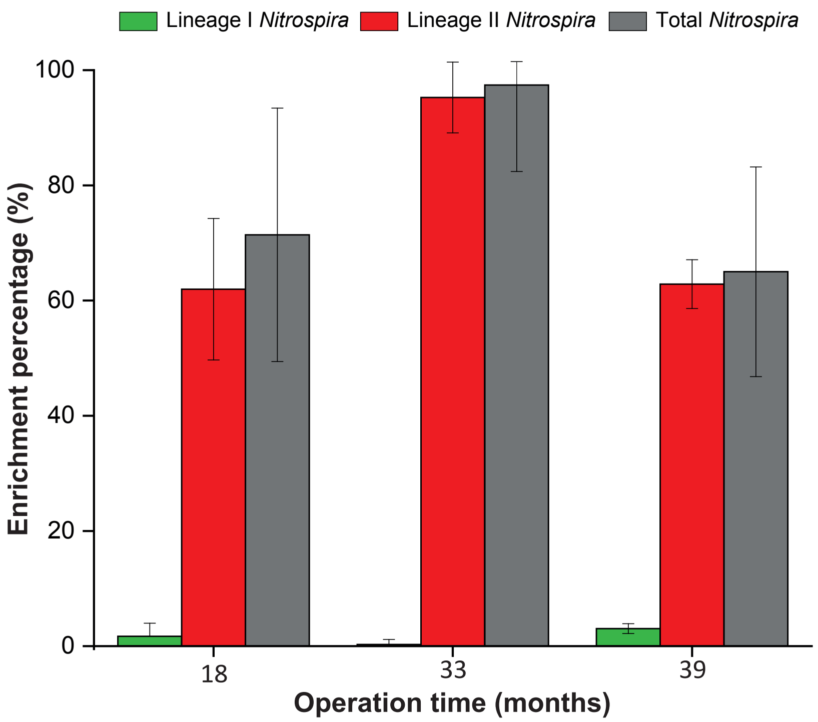


**Figure S2**. Relative abundance of sublineage I and II *Nitrospira* bacteria in the bioreactor system over the enrichment period. Sublineage abundances were normalized in relation to the relative abundance of the total *Nitrospira* population in the enrichment culture.

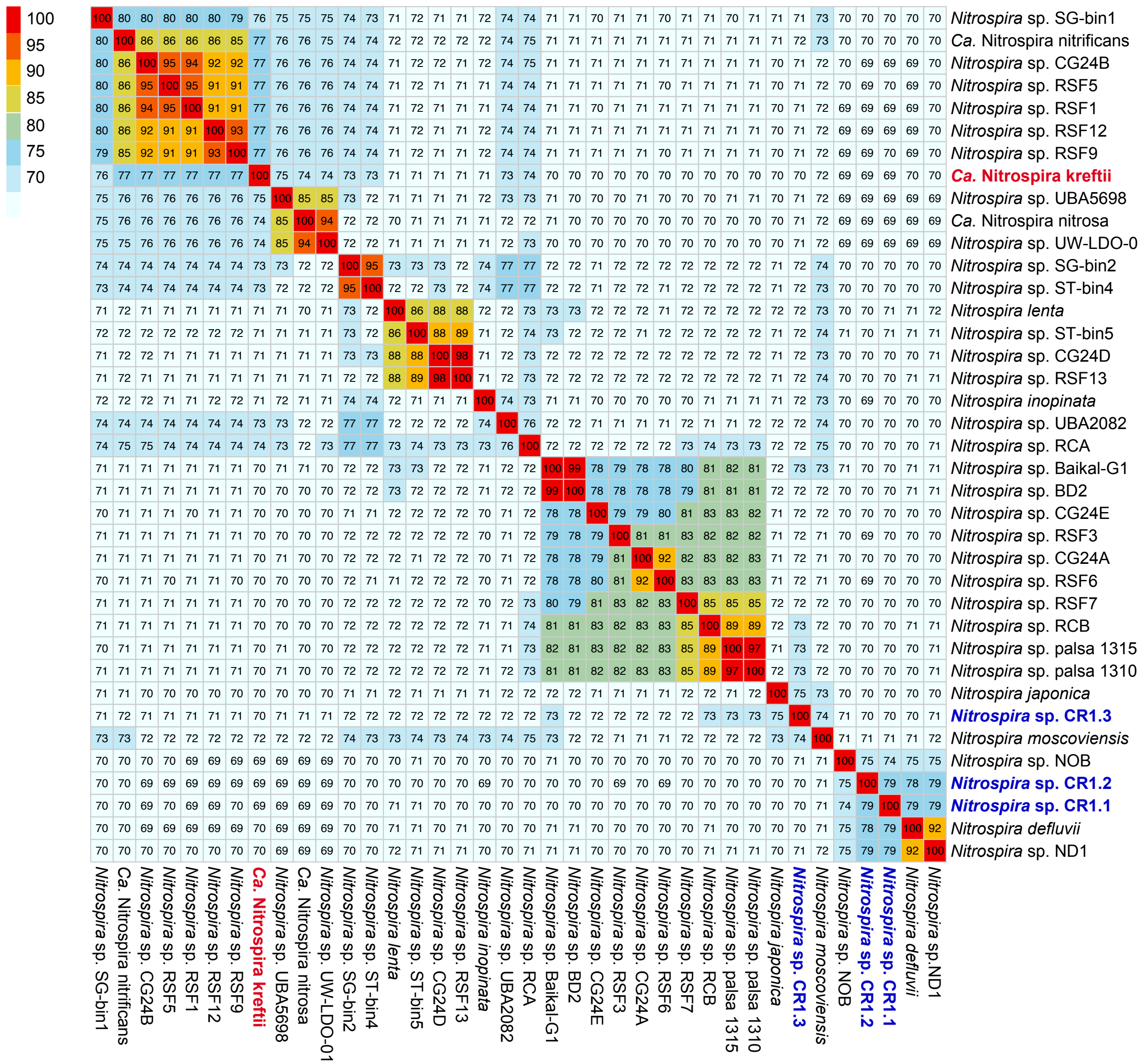


**Figure S3.** Genome similarity heatmap showing the pairwise ANI comparisons between the *Nitrospira* MAGs obtained in this study (in bold) and publicly available high-quality genomes of sublineage I and II *Nitrospira* species.

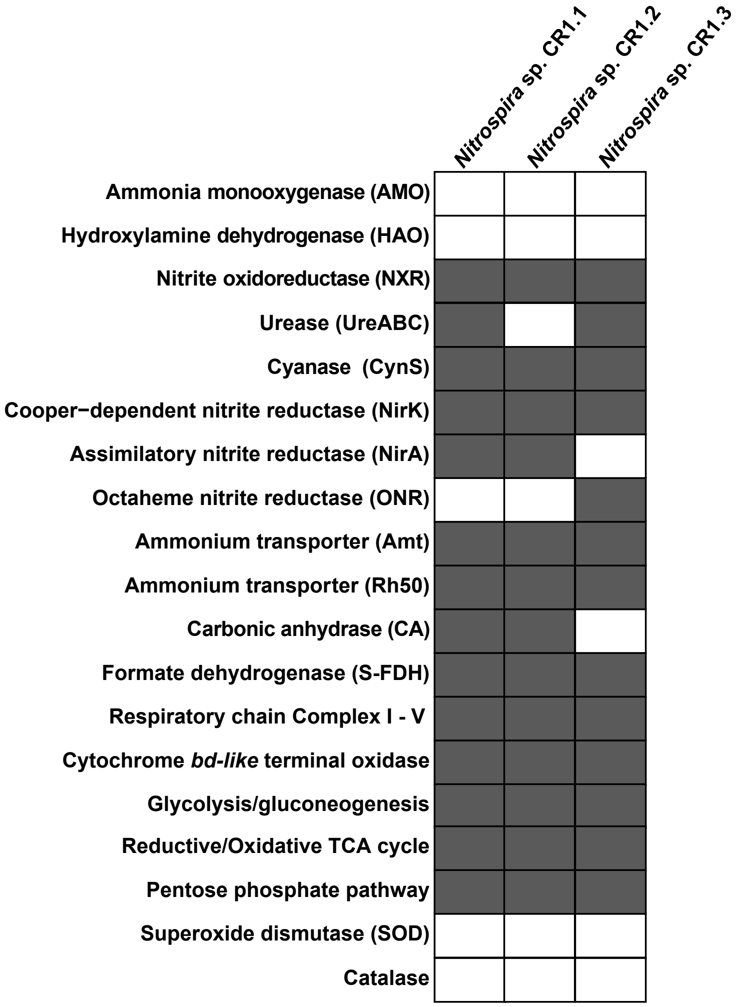


**Figure S4.** Distribution pattern of key metabolic features involved in nitrogen and alternative energy metabolisms in the canonical nitrite-oxidizing *Nitrospira* MAGs retrieved in this study. Dark grey and white indicate presence and absence of the respective genes, respectively.

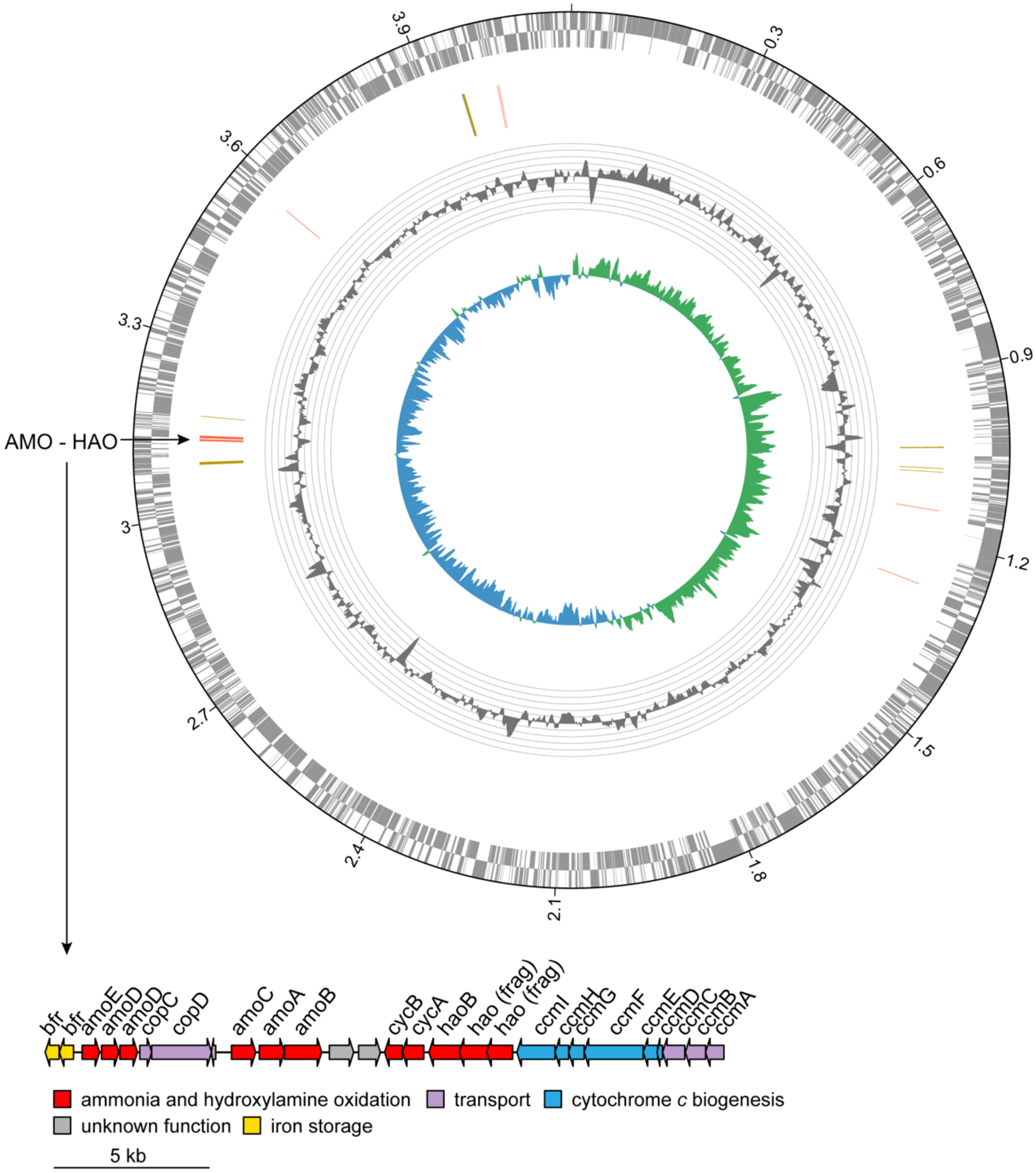


**Figure S5.** Circular representation of the "*Ca.* N. kreftii" chromosome. From outside to inside the rings display: (1) and (2) predicted coding sequences on forward and reverse strand, respectively, (3) Genes involved in ammonia and nitrite oxidation. Red: *amoABC*, *haoAB* and *cycAB*; orange: *nxrABC*. (4) Local GC bias and (5) GC skew (green: positive, blue: negative).

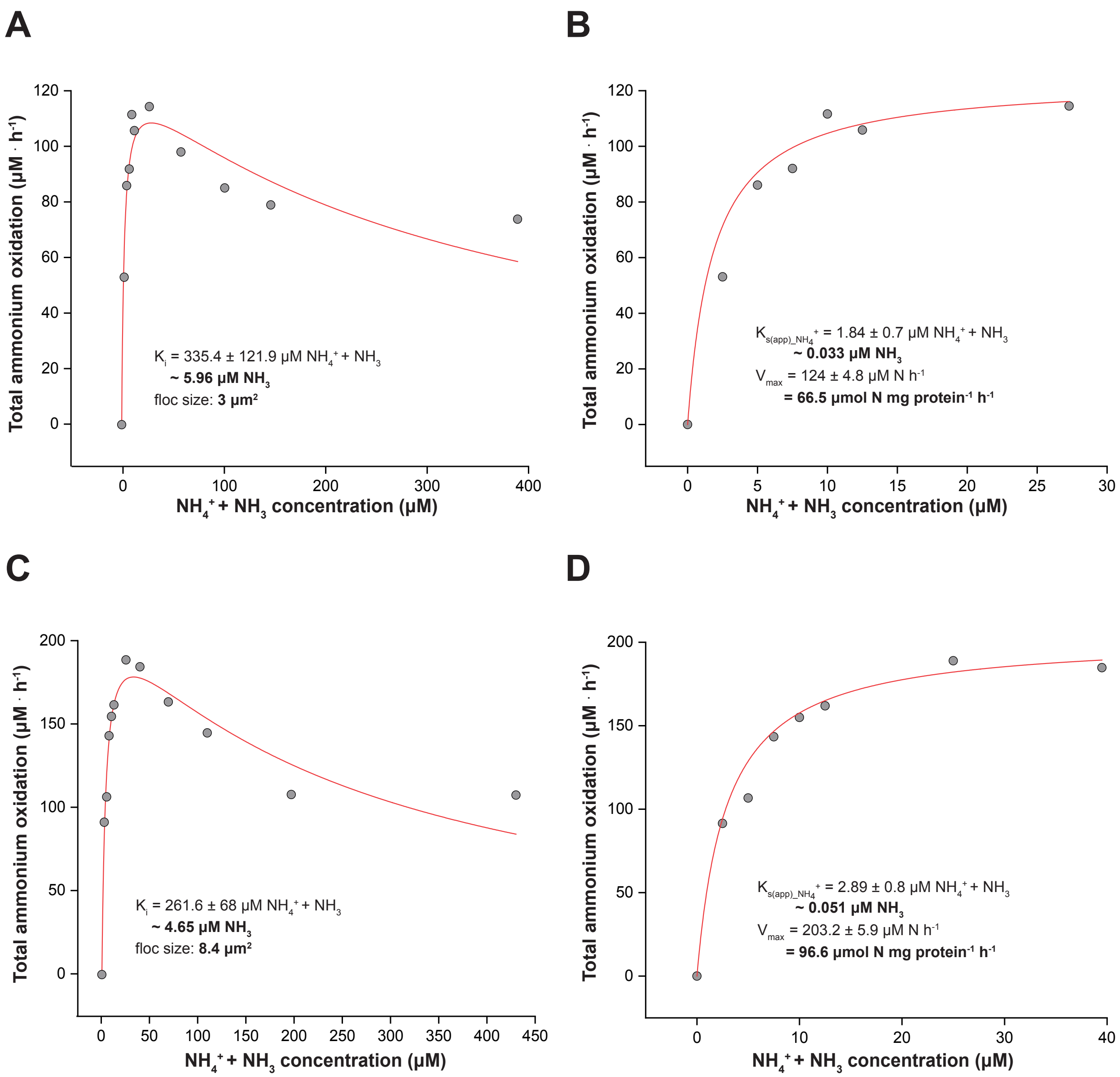


**Figure S6.** Ammonium oxidation kinetics of the "*Ca.* N. kreftii" enrichment culture. The red curves indicate the best fit (A, C) of all data to the substrate inhibition model and (B, D) of the data retrieved for non-inhibitory ammonium concentrations in a Michaelis-Menten kinetic equation. The reported standard errors are based on nonlinear regression. Results for two biological replicates are shown here; a third biological replicate is shown in Figure 4.

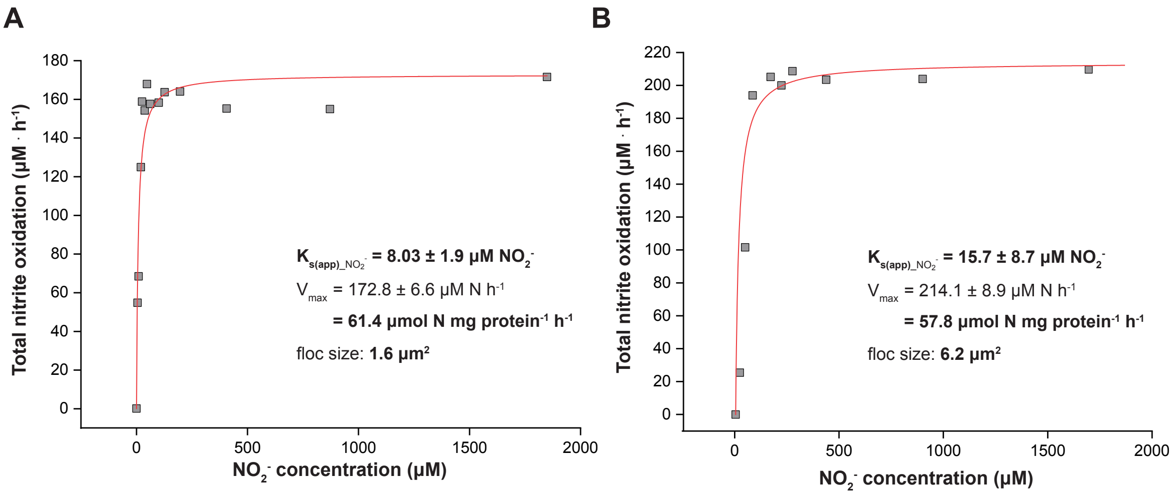


**Figure S7.** Nitrite oxidation kinetics of the "*Ca.* N. kreftii" enrichment culture. The red curves indicate the best fit of the data to the Michaelis-Menten kinetic equation. The reported standard errors are based on nonlinear regression. (A, B) Results for two biological replicates are shown here; a third biological replicate is shown in Figure 5.

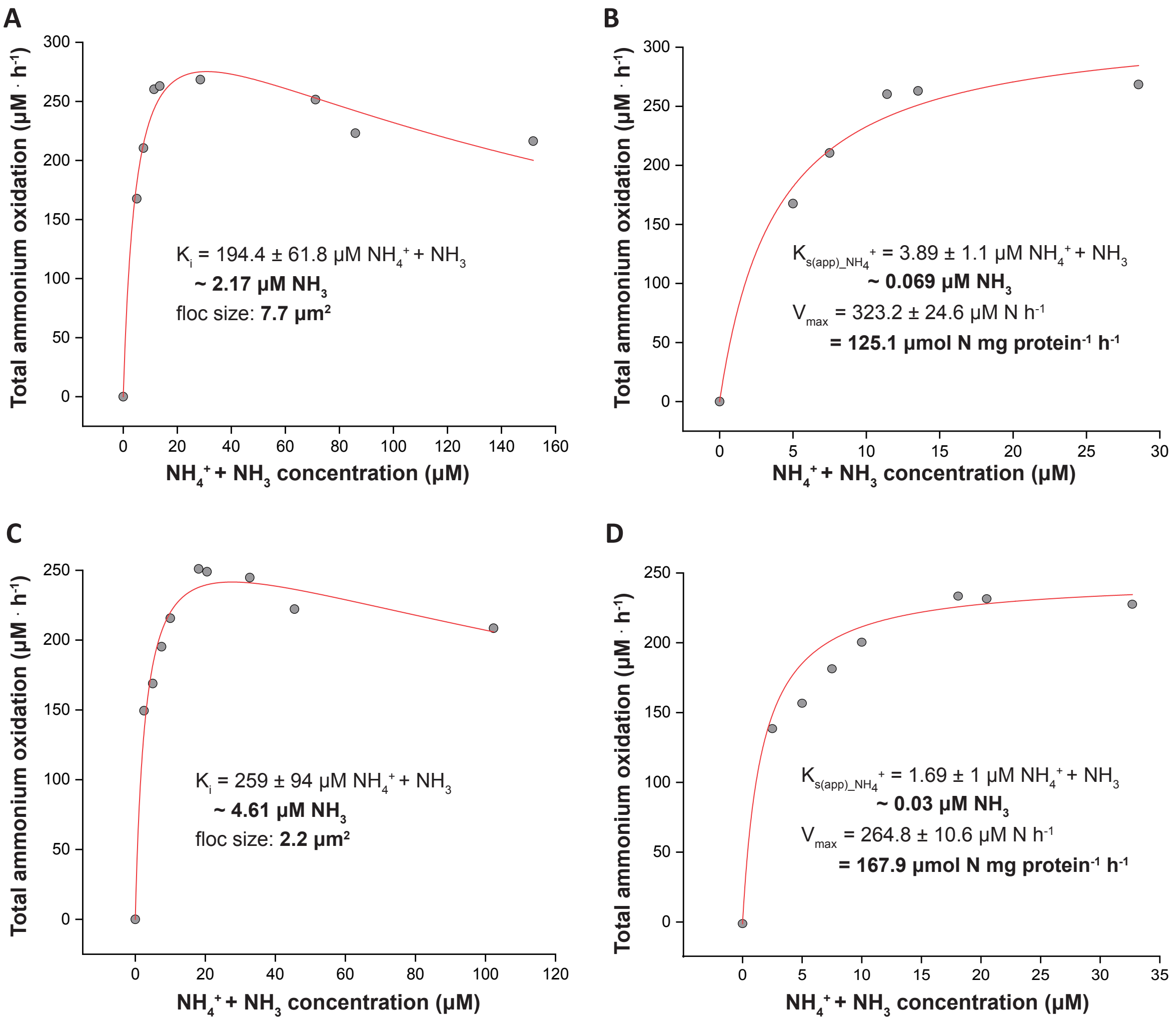


**Figure S8.** Ammonium oxidation kinetics of the "*Ca.* N. kreftii" enrichment culture, adapted to 1mM ammonium feeding. The red curves indicate the best fit (A, C) of all data to the substrate inhibition model and (B, D) of the data retrieved for non-inhibitory ammonium concentrations in a Michaelis-Menten kinetic equation. The reported standard errors are based on nonlinear regression. Results for two biological replicates are shown here; a third biological replicate is shown in Figure 6A, B.

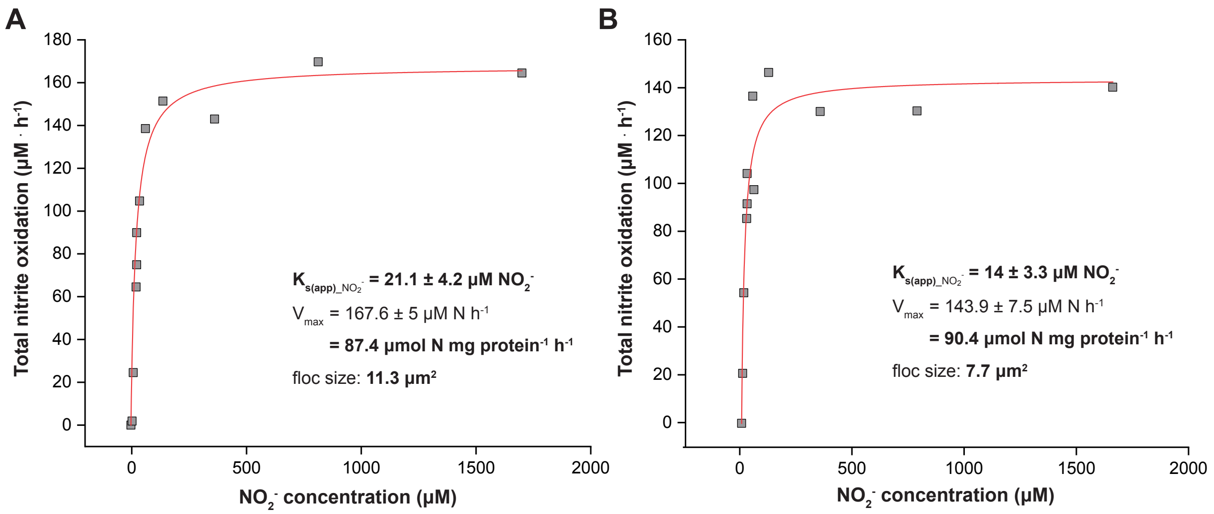


**Figure S9.** Nitrite oxidation kinetics of the "*Ca.* N. kreftii" enrichment culture, adapted to 1mM ammonium feeding. The red curves indicate the best fit of the data to the Michaelis-Menten kinetic equation. The reported standard errors are based on nonlinear regression. (A, B) Results for two biological replicates are shown here; a third biological replicate is shown in Figure 6C.

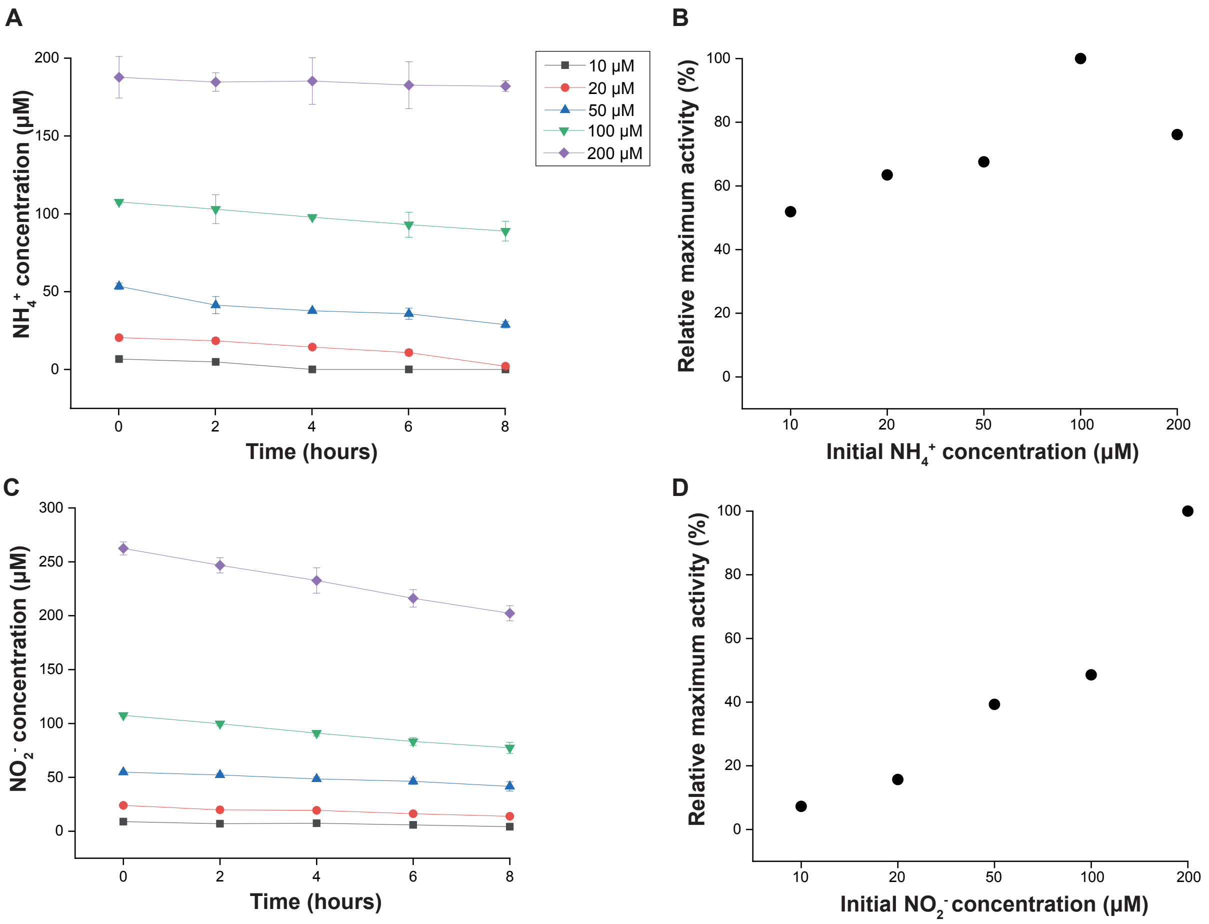


**Figure S10.** Oxidation of increasing concentrations of (A) ammonium and (B) nitrite in batch incubations inoculated with the "*Ca*. N. kreftii" enrichment culture. Relative maximum (C) ammonium and (D) nitrite oxidation rates were recorded at the substrate concentrations shown in (A) and (B) and calculated in relation to the maximum oxidation rate observed during the respective experiment. Symbols and error bars represent averages and standard deviations of three biological replicates, respectively.

**Table S1.** Specifications of the FISH probes used in this study.

| Probe | Target | FA %[a] | Sequence (5’→3’) | Reference |
|---|---|---|---|---|
| **Ntspa712** | *Nitrospira* phylum (most members) | 35 | CGCCTTCGCCACCGGCCTTCC | (1) |
| **Comp-Ntspa712** | Competitor to Ntspa712 | - | CGCCTTCGCCACCGGTGTTCC | (1) |
| **Ntspa662** | genus *Nitrospira* | 35 | GGAATTCCGCGCTCCTCT | (1) |
| **Comp-Ntspa662** | Competitor to Ntspa662 | - | GGAATTCCGCTCTCCTCT | (1) |
| **Ntspa1431** | Sublineage I of the genus *Nitrospira* | 35 | TTGGCTTGGGCGACTTCA | (2) |
| **Ntspa1151** | Sublineage II of the genus *Nitrospira* | 35-40 | TTCTCCTGGGCAGTCTCT CC | (2) |
| **EUB338 (Bact338)** | Most bacteria | 0-50 | GCTGCCTCCCGTAGGAGT | (3) |
| **EUB338 II (SBACT P 338)** | *Planctomycetales* | 0-50 | GCAGCCACCCGTAGGTGT | (4) |
| **EUB338 III (SBACT V 338)** | *Verrucomicrobiales* | 0-50 | GCTGCCACCCGTAGGTGT | (4) |

[a]Concentration of formamide (FA) in the hybridization buffer

**Table S2.** Enrichment of the bioreactor's biomass in *Nitrospira* bacteria over the total enrichment period determined by quantitative FISH.

| Operation time (months) | Enrichment (%) | S.E. (±) |
|---|---|---|
| 0 | 5.4 | 8.9 |
| 8 | 53 | 17.9 |
| 11 | 71.3 | 25.5 |
| 14 | 73.7 | 14.7 |
| 17 | 57 | 29.5 |
| 18 | 64.8 | 15 |
| 19 | 83.5 | 14.9 |
| 24 | 65.3 | 18.9 |
| 26 | 85.1 | 6.5 |
| 27 | 90.5 | 7.6 |
| 28 | 86.1 | 10 |
| 33 | 71.7 | 8 |
| 35 | 55.9 | 17.5 |
| 39 | 71.9 | 18.2 |

**Dataset S1 (separate file).** Overview of the medium- and high-quality metagenome-assembled genomes (MAGs; completeness ≥75%, contamination ≤10%) obtained from the enrichment culture after 17 months of enrichment.

**Dataset S2 (separate file).** Overview of the medium- and high-quality metagenome-assembled genomes (MAGs; completeness ≥75%, contamination ≤10%) obtained from the enrichment culture after 39 months of enrichment.

**Dataset S3 (separate file).** "*Ca.* Nitrospira kreftii" proteins with predicted functions in key metabolic pathways.
